# The primary motor cortex is critical for the retention of implicit sensorimotor adaptation

**DOI:** 10.1101/096404

**Authors:** Li-Ann Leow, Welber Marinovic, Timothy J Carroll, Stephan Riek

## Abstract

Sensorimotor adaptation, or adaptation of movements to external perturbations, is thought to involve the primary motor cortex (M1). In addition to implicit error-driven remapping, explicit re-aiming strategies also contribute to sensorimotor adaptation. However, no studies to date have examined the role of M1 in implicit learning in isolation from explicit strategies. Because the application of explicit strategies requires time, it is possible to emphasise implicit learning by controlling the time available to prepare movement. Here, we examined the role of M1’s role in implicit adaptation to rotated visual feedback whilst suppressing the use of explicit re-aiming strategies by limiting movement preparation times to less than 350ms. Perturbing M1 activity via single-pulse TMS during adaptation to a 30 ° rotation of visual feedback did not alter the rate or extent of error compensation, but elicited poorer retention in post-adaptation trials with no perturbation. This work shows that M1 is critical in the retention of new visuomotor maps as a result of implicit adaptation to a perturbation in sensory feedback when strategic error correction processes are suppressed.

**Highlights:**

- Adaptation of movements to perturbations occurs through explicit and implicit processes.
- Here, explicit strategies were suppressed by shortening movement preparation time.
- Perturbing motor cortex (M1) with TMS selectively impaired retention but not acquisition of sensorimotor adaptation.
- M1 plays a crucial role in retention of sensorimotor adaptation obtained via implicit learning.

---

Humans display remarkable flexibility in their ability to adapt to external perturbations, such as external force-fields (Shadmehr & Mussa-Ivaldi, 1994), or distortions of visual feedback (Welch, 1978; Cunningham, 1989). This form of learning, termed sensorimotor adaptation, is retained even up to a year (Shadmehr & Brashers-Krug, 1997; Landi *et al.*, 2011).

Converging evidence implicates a role for the primary motor cortex (M1) in the formation of motor memories during sensorimotor adaptation. For example, non-invasively altering M1 excitability alters the retention of learning in adaptation to force-field (Cothros *et al.*, 2006; Richardson *et al.*, 2006; Hunter *et al.*, 2009; Herzfeld *et al.*, 2014) and visuomotor rotation (Galea *et al.*, 2011; Riek *et al.*, 2012; Leow *et al.*, 2014; Villalta *et al.*, 2015). One recent study demonstrated that tagging M1 activity but not posterior parietal cortex activity with tDCS allows the concurrent learning of two opposite force-fields, which suggests that M1 is crucial to the formation of distinct motor memories. Similarly, neural recordings in M1 in non-human primates demonstrate that sensorimotor adaptation evokes a persistent change in the preferred direction of motor cortex neurons (Gandolfo *et al.*, 2000; Li *et al.*, 2001). This change in the preferred direction persists despite returning motor output to the unadapted state with no-perturbation trials (Mandelblat-Cerf *et al.*, 2011) and subsequent adaptation to the same perturbation (Paz *et al.*, 2003) or to an opposing perturbation (Zach *et al.*, 2012). How M1 plays a role in the encoding and retrieval of motor memories is less well understood, however, a recent study in non-human primates suggest that the motor cortices provide error signals that drive sensorimotor adaptation (Inoue *et al.*, 2016). Two key findings from this study support this proposal. First, neural recordings in the motor cortices showed greater M1 firing at a critical time window of 100 ms post-movement during sensorimotor adaptation, suggestive of a role of M1 in encoding errors post-movement. Second, when these M1 neurons were stimulated immediately after movement execution but not after this 100ms post-movement interval, errors during sensorimotor adaptation increased.

In humans, errors imposed by external perturbations can be reduced either through (1) explicit mechanisms, for example by strategically re-aiming clockwise to a target to compensate for a counterclockwise rotation in visual feedback (Uhlarik, 1973; Mazzoni & Krakauer, 2006; Benson *et al.*, 2011), or through (2) implicit mechanisms, where errors are incrementally reduced in an automatic fashion (Squire, 1992; Redding & Wallace, 1996). Explicit adaptive processes are used to correct errors to ensure success in the task, and their contribution to behavior is modulated to accommodate the current extent of implicit adaptation (Taylor *et al.*, 2014; McDougle *et al.*, 2015). It is therefore difficult to assess how a factor such as brain stimulation affects implicit adaptation from performance when explicit contributions are allowed. Thus, despite growing evidence demonstrating the role of M1 in the formation of motor memories during sensorimotor adaptation, it remains unclear whether M1 plays a role in the implicit and/or explicit mechanisms known to be active during sensorimotor adaptation (Taylor *et al.*, 2014). One possibility is that M1 is primarily involved in implicit mechanisms during sensorimotor adaptation. For example, M1 anodal tDCS only increased the persistence of adapted movements when there was no reason to recruit explicit mechanisms for error reduction (i.e., in when all feedback of the movement trajectory was hidden), but not when it is beneficial to use explicit mechanisms to reduce errors (i.e., when errors were revealed via feedback of the movement trajectory) (Galea *et al.*, 2011). Similarly, perturbing M1 activity reduced the persistence of adapted movements only when the perturbation was gradually imposed but not when the perturbation was abruptly imposed (Hadipour-Niktarash *et al.*, 2007). Gradual perturbations typically evoke small errors that often escape conscious awareness (Kagerer *et al.*, 1997), and thus are thought less likely to evoke explicit processes such as strategic re-aiming. The role of M1 in the rate or extent of implicit learning during visuomotor adaptation is not fully understood, as the majority of studies do not explicitly dissociate between implicit and explicit learning during sensorimotor adaptation. One way to isolate the contribution of implicit mechanisms in sensorimotor adaptation is to suppress the use of explicit mechanisms by reducing the amount of time available to prepare the movement (Haith *et al.*, 2015; Leow *et al.*, 2016). Although this procedure does not necessarily eliminate explicit processes when there is deliberate intention to re-aim, it appears sufficient to suppress the inclination to re-aim when targets are presented in a large angular range (Leow *et al.*, 2016), such that the rate of error compensation when movement preparation time is reduced is comparable to that of implicit learning estimated by subtracting self-reported aiming directions from actual movement directions (Taylor *et al.*, 2014). In this study, we examine the role of M1 in acquisition and retention of sensorimotor adaptation whilst suppressing the recruitment of explicit processes by reducing the amount of time available for movement preparation. We hypothesized that perturbing M1 excitability would affect retention but not the acquisition of sensorimotor adaptation when we suppressed the use of explicit strategies.

## Methods

Thirty-two right-handed participants (22 female, aged between 18–25 years old) with no recent wrist, elbow or shoulder injuries volunteered for the study. Right-handedness was confirmed with Edinburgh Handedness Inventory (Oldfield 1971). A medical questionnaire was used to screen the participants for neurological disorders and contraindications in relation to the application of TMS. The study was approved by the Medical Research Ethics Committee of The University of Queensland. All participants were briefed on the experimental procedures and gave written informed consent prior to the experiment, which conformed to the Declaration of Helsinki.

### Experimental design

Participants were randomly assigned into a ‘STIM’ or ‘SHAM’ groups in a counter-balanced manner. In each group, the participants performed at total of 520 trials of isometric force aiming tasks toward 8 radial targets on a computer screen located about 1 m away with their right hand. Their right forearms were secured into a custom-made manipulandum, described previously (de Rugy *et al.*, 2012), in the neutral wrist position (midway between pronation and supination). Both elbows were kept at 110° with the forearm parallel to the table and supported by the manipulandum. The wrists were fixed by a series of twelve adjustable metal clamps contoured around the metacarpal-phalangeal joints and around the wrist proximal to the radial head. Wrist forces in abduction-adduction and flexion-extension directions were recorded via a six degree-of-freedom force transducer (JR3 45E15A-163-A400N60S, Woodland, CA) attached to each manipulandum. Force data were sampled at a rate of 2 kHz via two 16-bit National Instruments A/D boards (NI BNC2090A, NI USB6221, National Instruments Corporation, USA). The forces exerted in flexion-extension and abduction-adduction directions were displayed as a cursor that moved in two dimensions (x = flexion-extension, y = abduction-adduction) on the computer screen via a custom written Matlab program. During the experiment, participants received 240 stimuli at the left motor cortex for the ‘STIM’ group or at the vertex for the ‘SHAM’ group out of the 520 trials of isometric force aiming tasks. Stimulation onset was timed to when the movement extent reached the distance between the start and the target, similar to previous work where TMS onset was timed to the trial end (Hadipour-Niktarash *et al.*, 2007)

### Isometric force aiming tasks

Participants were required to move a cursor that represented the resultant force exerted at the wrist joint in 2 degrees of freedom (ab-adduction versus flexion-extension) to 8 radial targets which were 45° apart for 520 trials. Target location was randomized. Before starting the experiment, participants completed at least 16 trials (2 trials per target) to familiarize themselves with the target-aiming task. The experiment consisted of 4 blocks, which proceeded as follows. (1) **Pre**-adaptation: 160 trials with no rotation of visual feedback. (2) **Adaptation**: 240 trials with a 30° of rotation of visual feedback of the cursor position, either clockwise or counter-clockwise relative to the center of the start circle. For example, when an upward movement was made toward a target with radial deviation in the wrist, participants in the clockwise rotation group experienced a rightward deviation of the visual feedback of the cursor for a clockwise 30° visual rotation. Likewise, participants in the counterclockwise rotation group experienced a leftward deviation of the cursor for a counter-clockwise 30° visual rotation for the same upward target. (3) **No-feedback**: 40 trials with no visual feedback of cursor position, and no rotation of visual feedback—unlike in our recent work (Leow *et al.*, 2016), participants were not informed that the rotation was removed, in order to ensure close adherence to the previous M1 TMS study (Hadipour-Niktarash *et al.*, 2007), and (4) **Post**-adaptation: visual feedback of cursor position with no rotation.

Each trial began with a visual word cue “Relax” to prompt participants to relax their right forearm. A cursor corresponding to their wrist forces was displayed within a red circle indicating the origin of the computer screen. The participants were provided with 4 beats of audio cue and required to initiate cursor movement toward the target as quickly as possible on the 4^th^ beat. The targets appeared as green circles at 75 % of the distance from the origin to the edge of the computer screen (10 cm) at 300 ms prior to the 4^th^ audio beat. Participants were given post-trial feedback on whether the movements were too early (<150ms) or too late (>350ms) to encourage consistent movement onsets and, as a result, consistent preparation time. We selected 350ms as a cutoff because piloting showed that it was very difficult to move accurately to the target using the wrist manipulandum at briefer cutoff times. The cursor gain was set such that 20 N was required to reach the edge of the screen. The 8 targets appeared in a random order within every cycle of 8 trials to prevent participants from anticipating the location of upcoming target.

### Surface electromyography recordings

Electromyography (EMG) signals were recorded from the flexor carpi radialis (FCR) and extensor carpi radialis brevis (ECRb) of the right forearm. Standard skin preparation was performed after the muscles were located and marked. Bipolar Ag/AgCl surface electrodes placed on the belly of the forearm muscles with an inter-electrode distance of 2 cm (centre to centre). The EMG signals were amplified with a gain of 500 ~ 1000 with Grass P511 amplifiers (Grass Instruments, AstroMed, West Warwick, RI) and band-pass filtered (10 Hz - 1 kHz).

### Transcranial magnetic stimulation

Single-pulse TMS was used to elicit motor evoked potential (MEP) responses from the FCR and ECR muscles via a 70 mm outer diameter figure-of-eight magnetic coil (Magstim 200, Magstim, UK) over the forearm area of the left motor cortex. The magnetic coil was held tangentially on the scalp with the handle pointing backwards and 45° away from mid-sagittal axis inducing a current in the posterior-anterior direction. An optimal location that elicited the largest and most consistent MEP response was determined and marked with felt-tip pen to ensure consistent stimulation throughout the experiment. The resting motor threshold (RMT) for each participant was determined as the minimum stimulus intensity that produced a MEP response of approximately 50 μν in 5 out of 10 consecutive trials while the right forearm was at rest in the manipulandum. The testing intensity was set to 120% of each participant’s RMT.

### Data analysis

Data reduction was performed using custom Matlab software (Mathworks). Forces in *x* and *y* axes were transformed to screen coordinates (e.g. cursor position) and filtered using a low-pass 2^nd^ order Butterworth filter with a cut-off frequency of 10Hz. Movement onsets were estimated from the tangential speed time series (Teasdale *et al.*, 1993). Movement direction was computed as the angle between the initial position of the cursor at movement onset and its position 100 ms later, which is sufficient to prevent the use of online feedback mechanisms to correct cursor trajectory (Desmurget & Grafton, 2000).

Statistical analyses were run on cycle-averaged data, with 1 cycle defined as 8 trials. The dependent variable of interest was directional error, which was defined as the difference between movement direction and target direction. Intersubject differences in baseline directional biases (Ghilardi *et al.*, 1995), are known to affect adaptation behaviour [36–38]. Directional errors were bias-corrected by subtracting directional errors from the mean of the first 6 cycles (we did not bias correct using the last baseline cycle (Cycle 20) here, because previous research has shown that directional biases reduce with training. Unlike previous studies which had fewer baseline cycles than in this study(Taylor *et al.*, 2014). Qualitatively similar results were obtained when we estimated baseline biases from baseline cycle 6, which is what we did in our previous work (Leow *et al.*, 2016). In the adaptation, no-feedback, and washout blocks, only movements that were within 120° of the target (i.e., 60° clockwise or counterclockwise of the target) were included for statistical analyses. This outlier removal procedure excluded less than 2.5% of the data for each phase. Stimulation (sham, stimulation) x Rotation Direction (CCW, CW) x Cycle ANOVAs were run on each block. For all ANOVAs, we used generalized eta-squares (rather than partial eta-squares) to estimate effect sizes. Eta-squares between 0.01 to 0.06 were considered small, eta-squares greater than 0.06 but smaller than 0.14 were considered medium, and eta-squares in excess of 0.14 were considered large (Cohen, 1988).

To examine the rate of adaptation without the possible confound of intrinsic bias in movement direction (Hardwick & Celnik, 2014), we also fit cycle-averaged movement directions for each dataset to a single-rate exponential function (Zarahn *et al.*, 2008), as follows:

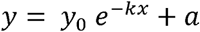

where y is the movement direction, x is the trial number, k is the rate constant that indicates the rate with which movement direction changes, a is the movement direction at which performance reaches asymptote, and y0 + a is the hypothetical y value when x is zero. One dataset failed to fit to the curve. Independent samples t-tests were used to compare rate constants from the sham and the stimulation group.

We also examined the rate of decay in the no-feedback block by fitting trial-by-trial movement directions for the no-feedback block to a straight line, as follows:

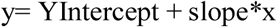

where y is the movement direction, x is the trial number, slope is the rate constant that indicates the rate with which movement direction changes, and YIntercept is the hypothetical y value when x is zero. We did not use cycle-averaged movement directions for the no-feedback block as there were only 5 no-feedback cycles, and this was insufficient data to fit to the straight line. Independent samples t-tests were used to compare slopes from the sham and the stimulation group.

To additionally guard against the possibility that individual differences in intrinsic movement biases influenced statistical comparisons between the stim and the sham TMS groups, we also quantified adaptation behaviour as a change in performance within each block. For directional error data, this was quantified as the difference in directional errors from the first cycle of each block to the last cycle of each block.

To test whether participants complied to the preparation time constraints throughout learning, we evaluated temporal error for the preparation time, defined as the error in time between target appearance and the time of movement onset. To guard against the possibility that any differences between the stim and the sham TMS groups resulted from individual differences in baseline ability to adhere to the preparation time constraints, we also quantified the change in preparation time within each block as the difference in preparation time from the first cycle to the last cycle of each block. Group differences on these change scores were tested using unequal variance t-tests (Welch’s t-test) to compare changes in directional error and preparation time across all phases of the experiment. The 95% confidence intervals (CI) were used for pairwise differences.

## Results

We first used t-tests to verify whether initial performance between groups differed in terms of directional error or preparation time at the beginning of each phase of the experiment. We found no differences in terms of mean directional error (all Ps > 0.38) or preparation time (all Ps > 0.13) across all experimental phases. These results indicate the sham and the stim groups did not differ reliably in initial performance at the beginning of the experimental phases.

Figure 1 shows cycle-averaged directional errors for each phase (1 cycle=8 trials). Each block was split into 2 sub-blocks for analysis (early (cycles 1–15) and late (cycles 1530). Stimulation (sham, stimulation) x Rotation Direction (CCW, CW) x Cycle x Phase (early, late) ANOVAs were run. There was no significant main effect of stimulation, and stimulation did not interact significantly with any other factor (all p>0.3). Analyses of rate constants obtained from fitting the single-exponential function to individual datasets yielded similar results (see Figure 1B)—sham and stim M1 TMS groups did not differ reliably in rate of error reduction in the adaptation phase t(29) =0.17, p=.86, cohen’s d=.06, small effect size).

**Figure 1.**
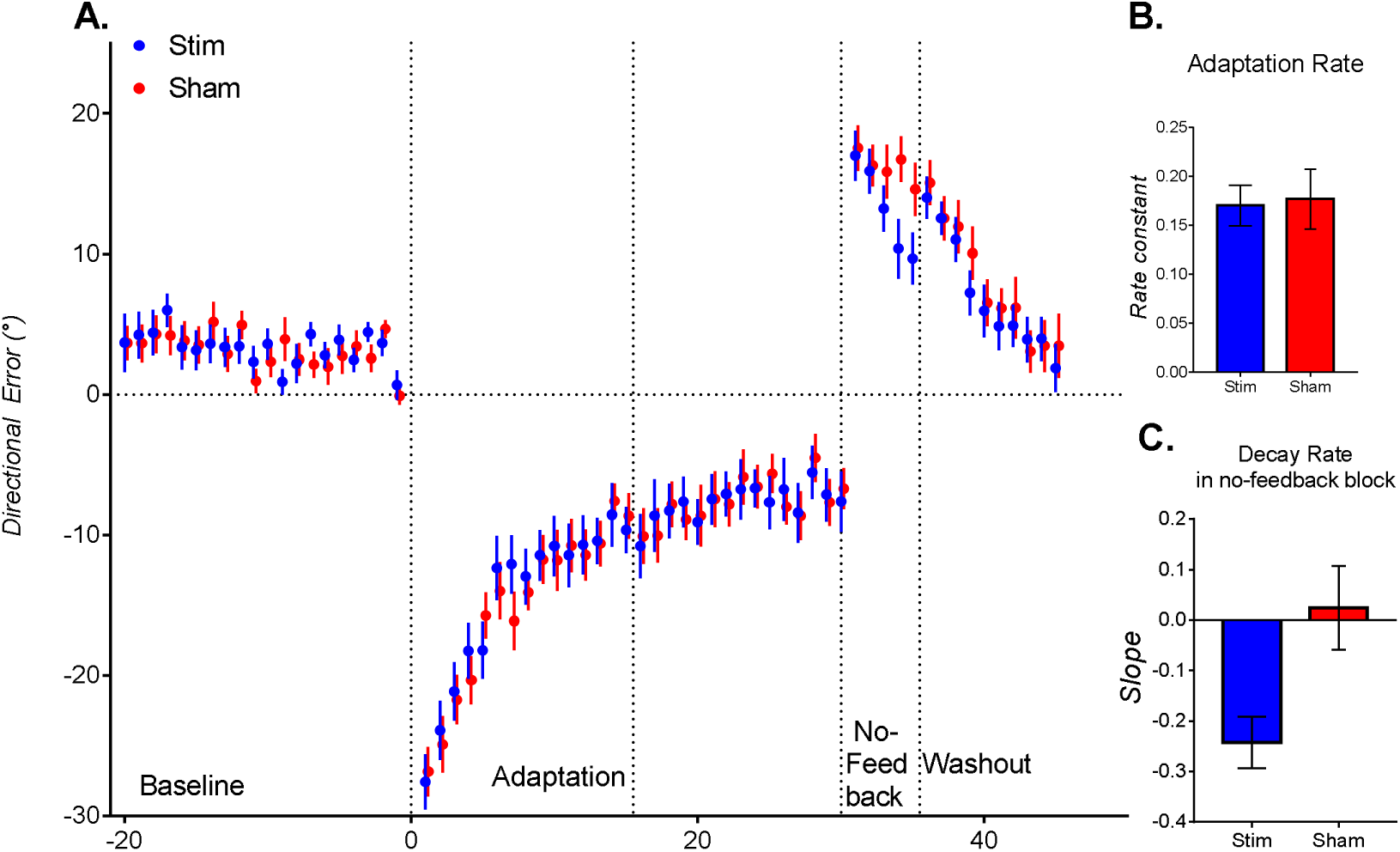
**A**: Group mean cycle-averaged directional errors for the group who encountered sham TMS pulses (1 cycle= 8 trials). Error bars indicate standard errors of the mean. Data from the clockwise rotation group were sign-transformed to allow statistical comparisons between clockwise and counterclockwise groups. Data from the adaptation phase, no-feedback phase, and the washout phase were bias corrected by subtracting estimated intrinsic bias (i.e., mean movement direction from the first 6 baseline cycles). Note that for the no-feedback block, greater decay was evident in the STIM group who received single-pulse TMS on the estimated location of M1 during adaptation than the SHAM group who received TMS on the vertex. **B**: Adaptation rate, as estimated by rate constants fit to cycle-averaged individual data from the adaptation block. **C**. Decay rate of trial-by-trial errors from the no feedback block for the sham group (blue bar) and the stim group (red bar). Error bars indicate standard errors of the mean.

In the first cycle of the no-feedback phase, movements remained in the adapted state, resulting in large errors in the opposite direction to the rotation indicating retention of the learning. Consistent with previous findings(Kitago *et al.*, 2013), despite the absence of visual feedback of the movement, and despite the experimenter not providing explicit knowledge that the rotation had been removed, movements decayed with increasing cycles, as there was a significant main effect of cycle F(4,112)=6.60, p <0.001, eta-squared = .17, as movements decayed across cycles. M1 stimulation resulted in poorer retention in this no-feedback block, as shown in faster decay of movements to the unadapted state in the retention block (see Figure 1C), as a Stimulation (Sham, Stim) by Cycle (1–5) by Rotation Direction (CW, CCW) ANOVA revealed a significant Stimulation x Cycle interaction. F(4, 112)=2.79, p = .03, eta-square =.09 (moderate effect size). The rate of decay in the no-feedback phase (as shown by slopes fit to trial-by-trial data) was faster for the stimulation group than the sham stimulation group, as slopes were significantly more negative in the stimulation group than the sham stimulation group, t(30) = 2.83, p =.005, cohen’s d= 0.96 (large effect size). After the no-feedback phase, visual feedback was returned in the post-adaptation phase.

Close inspection of Figure 1A suggests that unlike participants who received sham TMS, participants who received M1 TMS showed an apparent rebound in errors upon return of visual feedback in the washout phase (i.e., despite the greater decay of adaptation in the no-feedback block, they showed a return to movements that were more adapted to the rotation). One possibility is that the return of visual feedback acted as a contextual cue for retrieval of the motor memory that transiently decayed as a result of disrupting M1 activity. To test this, we ran a Stimulation (Sham, Stim) by Cycle (Last No Feedback cycle, First Washout Cycle) by Rotation Direction (CW, CCW) ANOVA. However, the Cycle x Stimulation interaction was not reliable, F(1, 28)=2.67, p =.11, although this was a moderate effect size (eta-squared =.07), and the main effect of stimulation was also not reliable (p=.38), To evaluate the effect of stimulation for the washout block, we ran a Stimulation (Sham, Stim) by Cycle (1–10) by Rotation Direction (CW, CCW) ANOVA. The main effect of stimulation was not reliable, and did not interact reliably with any factor (all p>0.3s).

As shown in Figure 2A, the analysis of change in directional error detected a statistically reliable difference between groups in the Retention block (t_29.96_ = 2.57, p = 0.015, 95% CI [−1.49, 12.87]), indicating that the group receiving TMS stimulation retained less of the original adaptation without visual feedback of the cursor position. T-tests failed to detect statistically reliable differences between groups in all other phases of the experiment (Pre: t_26.74_ = 0.34, p = 0.73, 95% CI [−6.75, 9.43]; Adaptation: t_29.41_ = 0.51, p = 0.61, 95% CI [−4.77, 7.93]; Post: t_28.66_ = 0.49, p = 0.62, 95% CI [−5.17, 8.51]). These results suggest M1 disruption sped up the decay of adapted movements, suggesting that M1 impaired retention of sensorimotor adaptation.

**Figure 2.**
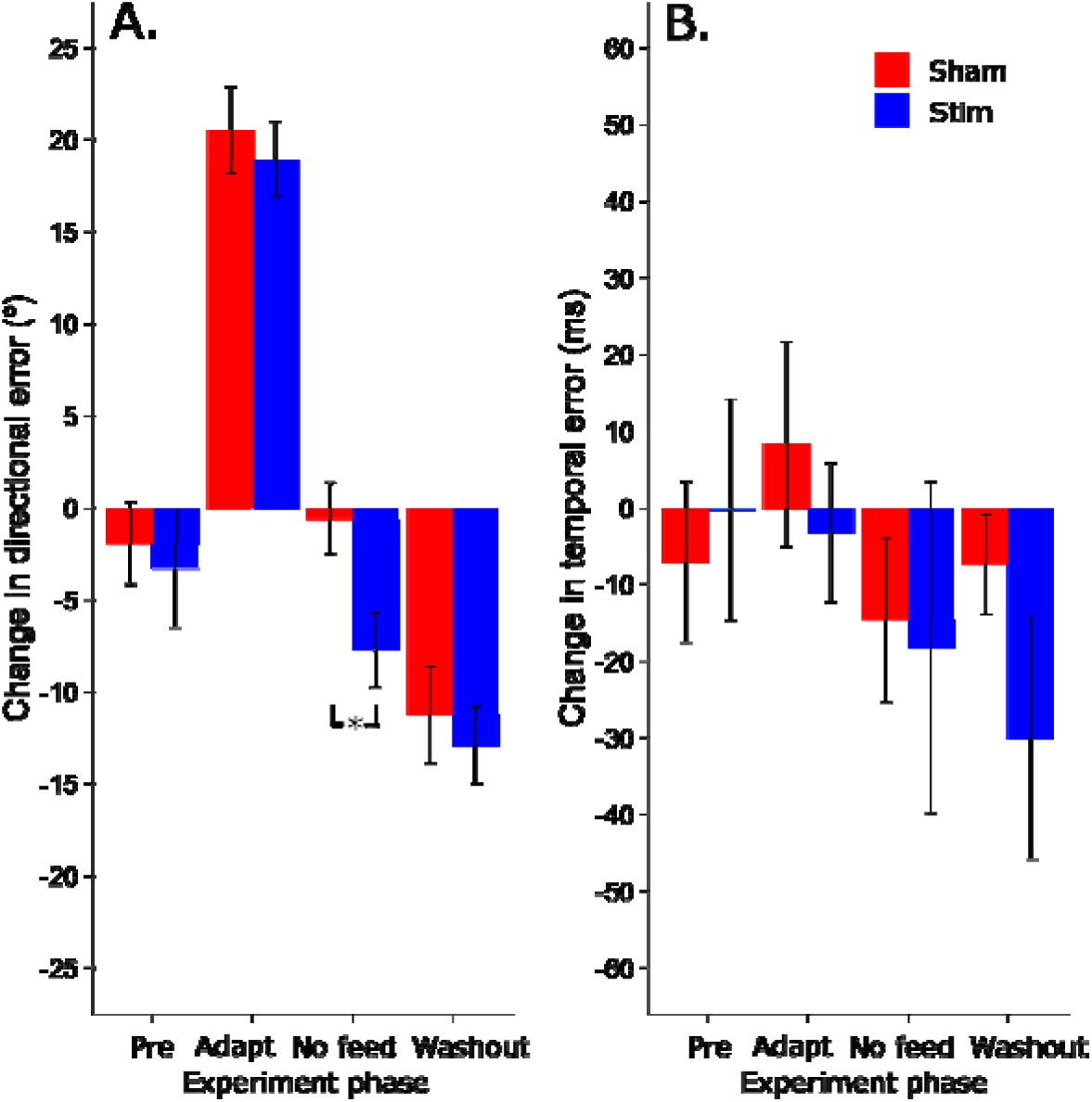
**A** - Mean changes in directional error (mean of first 4 trials – mean of last 4 trials) across all phases of the experiment. **2B** - Mean changes in temporal errors in preparation time (First 4 trials - Last 4 trials) across all phases of the experiment. *Mark statistically significant differences between groups. Error bars are 95% CI.

Figure 2B shows the changes in temporal errors for preparation time across all phases of the experiment. T-tests found no reliable differences between groups for any pairwise comparisons on preparation time (Pre: t_27.28_ = 0.37, p = 0.71, 95% CI [−43.38, 29.88]; Adaptation: t_26.39_ = 0.71, p = 0.48, 95% CI [−21.60, 44.69]; Retention: t_21.88_ = 0.14, p = 0.88, 95% CI [−46.46, 53.53]; Post: t_19.85_ = 1 32, p = 0.20, 95% CI [−13.04, 58.23]). These results indicate that any differences - or the lack thereof - in terms of directional error are unlikely to be due to differences in ability to adhere to the preparation time restraints.

## Discussion

Here, we show that when explicit re-aiming is likely suppressed by shortening movement preparation time, perturbing M1 during sensorimotor adaptation does not affect the rate and extent of error compensation, but elicits greater decay of adapted movements. This shows that M1 plays a key role in *retention* of sensorimotor adaptation acquired via implicit mechanisms. Although it is possible that constraining movement preparation time alone might not be sufficient to completely eliminate the employment of explicit strategies, our preparation time constraints appeared sufficient to reduce explicit strategy use to reveal implicit sensorimotor adaptation, in line with previous work showing that constraining preparation time results in slower error compensation typically indicative of reduced explicit strategy use, revealing implicit mechanisms in sensorimotor adaptation (Fernandez-Ruiz *et al.*, 2011; Haith *et al.*, 2015). This work adds to existing evidence (Hadipour-Niktarash *et al.*, 2007) supporting the hypothesis that M1 plays a role in implicit mechanisms of sensorimotor adaptation. In that study, M1 TMS impaired the retention of adapted movements with a gradually imposed perturbation, but not an abruptly imposed perturbation (note however that the stimulation protocol differed for the gradual and the abrupt perturbation—stimulation was applied in the pre-rotation phase for the abrupt condition but was applied in the rotation phase for the gradual perturbation) (Hadipour-Niktarash *et al.*, 2007). Such gradual perturbations evoke small errors which typically escape conscious awareness and thus are less likely to evoke strategic re-aiming. The abrupt perturbation was likely to have evoked explicit re-aiming, which should have sped up error compensation, and could have masked effects of M1 TMS on underlying implicit processes. However, the current data indicate that TMS to M1 does not impair acquisition of visuomotor adaptation even when explicit re-aiming is likely suppressed by constraining preparation time conditions.

It is worth noting that gradual and abrupt perturbations not only evoke differences in conscious awareness of the perturbation, but also differences in the amount of movement repetition of the adapted movement, as gradually imposed perturbations necessarily result in less repetition of the fully adapted movement than abrupt perturbations. One might argue that our findings of faster decay of adaptation learning when M1 activity was disrupted by TMS are driven by its effects on repetition-dependent effects (i.e., on use-dependent plasticity, where repetition of a movement induces a bias in subsequent movement toward that repeated movement) (Classen *et al.*, 1998). Previous studies have demonstrated that modulating M1 excitability selectively alters sensorimotor adaptation when repetition of the adapted movements are allowed. For example, one previous study (Orban de Xivry *et al.*, 2011) showed that the extent of error compensation was reduced by perturbing M1 excitability when the perturbation schedule afforded extended repetition of the adapted movement, but not when repetition was constrained. Another study showed that M1 excitability increased when an abrupt perturbation schedule encouraged extended repetition of the adapted movement, but not when a gradual perturbation schedule reduced repetition of the adapted movement [39]. Similarly, altering M1 excitability with tDCS only changed the persistence of adapted movements with extended repetition of the adapted movement, and not with limited repetition of the adapted movement (Leow *et al.*, 2014)—this effect was demonstrated when target manipulation was used to ensure extended repetition of the adapted movement in a gradual perturbation schedule, suggesting that the effect of M1 stimulation in that study was not due to the difference in the size of the error from abrupt and gradual perturbation schedules, but rather due to the amount of repetition of the adapted movement. However, although we cannot preclude the possibility that use-dependent plasticity effects contributed to our current results, we suggest that our results are not solely driven by use-dependent effects, as movements were made towards eight randomly presented targets, limiting continuous repetition of the adapted movement towards any one target.

We note that M1 TMS may also affect processing in remote brain regions (Bestmann *et al.*, 2004) known to be involved in sensorimotor adaptation such as the cerebellum, either directly via functional interconnections (e.g.,(Ugawa *et al.*, 1991)), or indirectly due to compensatory mechanisms (Lomber, 1999). However, given the weight of previous findings from multiple paradigms (brain stimulation, neural recordings, neuroimaging) (Gandolfo *et al.*, 2000; Li *et al.*, 2001; Paz *et al.*, 2003; Cothros *et al.*, 2006; Richardson *et al.*, 2006; Hunter *et al.*, 2009; Galea *et al.*, 2011; Mandelblat-Cerf *et al.*, 2011; Riek *et al.*, 2012; Zach *et al.*, 2012; Herzfeld *et al.*, 2014; Leow *et al.*, 2014; Panouilleres *et al.*, 2015; Villalta *et al.*, 2015) suggesting involvement of M1 in formation of motor memories in sensorimotor adaptation, we think that the most parsimonious account of our findings is that M1 is involved in retention of sensorimotor adaptation, even when explicit processes were suppressed with short preparation times.

Finally, it is important to note that our findings that M1 contributes to implicit learning do not preclude the possibility that M1 also contributes to explicit learning processes. For example, in humans, increasing M1 excitability improves error compensation when using large perturbations (60°) which engage explicit processes (Panouilleres *et al.*, 2015). Similarly, reducing M1 excitability also slowed reaction times when participants were reducing large errors and impaired error compensation when relearning the same perturbation (i.e., impaired savings) (Riek *et al.*, 2012). Several studies have shown that altering M1 excitability alters savings (Richardson *et al.*, 2006; Panouilleres *et al.*, 2015), which has recently been proposed to be at least partly driven by explicit mechanisms (Haith *et al.*, 2015; Morehead *et al.*, 2015).

